# Use of selenocysteine, the 21st amino acid, in the fungal kingdom

**DOI:** 10.1101/314781

**Authors:** Marco Mariotti, Gustavo Salinas, Toni Gabaldón, Vadim N. Gladyshev

## Abstract

Selenoproteins are a diverse class of proteins containing selenocysteine (Sec), the 21st amino acid, incorporated into proteins during translation through a unique recoding mechanism. Selenoproteins fulfil essential roles in several lineages, including vertebrates; yet, they are not ubiquitous across the tree of life. In particular, the fungal kingdom was thought to lack selenoproteins, a paradigm that we defy here. We show that the genetic machinery for Sec utilization is present in the genomes of five species belonging to diverse early-branching fungal phyla (Chytridiomycota, Zoopagomycota, and Mucoromycota). We identified several selenoprotein genes in each of these genomes, and we detected canonical Sec insertion RNA structures (SECIS elements) for some of them. The identified selenoproteins are known or predicted oxidoreductases, some of which are conserved in mammals. Phylogenetic analyses support a scenario of vertical inheritance of the Sec trait within eukaryotes and fungi. Sec was then lost in numerous independent events in various fungal lineages, even within Sec-containing phyla. Notably, Sec was lost at the base of Dikarya, resulting in the absence of this trait in Saccharomyces cerevisiae and other well studied fungi. Our results indicate that, despite scattered occurrence, selenoproteins are found in all kingdoms of life.

## INTRODUCTION

Selenoproteins are a diverse class of proteins that contain selenocysteine (Sec), known as the 21st amino acid in the genetic code. Sec is co-translationally inserted into proteins in response to UGA codon through an unusual recoding mechanism, wherein UGA, normally read as stop signal, is translated as Sec (1). Sec insertion occurs only in selenoprotein genes, due to the presence of RNA structures known as SECIS (SEC Insertion Sequence) elements (2, 3). A set of factors, here collectively referred to as Sec machinery, are dedicated to the synthesis and insertion of Sec. In eukaryotes, they include tRNASec (Sec-specific tRNA, acting as scaffold for Sec synthesis), SPS (selenophosphate synthetase), PSTK (phosphoseryl-tRNASec kinase) and SecS (Sec synthase) for the synthesis of Sec; and EFsec (Sec-specific elongation factor) and SBP2 (responsible for recognizing SECISes) for its insertion ((1, 4, 5) and references therein). Several additional proteins have been implicated in the Sec pathway (e.g. L30, nucleolin, Ser-tRNA synthetase), but in contrast to the core Sec machinery, they are not exclusively involved in the Sec trait.

Sec is typically found in the catalytic sites of oxidoreductase enzymes, several of which perform essential functions in mammals (e.g. redox homeostasis, hormone maturation) (1). For most selenoproteins, homologs exist that replace Sec with Cys. It is believed that Sec confers a selective advantage over Cys in proteins, which is generally attributed to enhanced reactivity, and/or increased resistance to irreversible oxidative inactivation (6, 7). Mammals possess 24-25 selenoprotein genes (8, 9), which are highly conserved and resistant to both gene loss and Sec-to-Cys conversion events (10).

However, selenoproteins are not present in all organisms, as further testified by the lack of Sec machinery in their genomes. Bacteria show a scattered distribution of the Sec trait, encompassing approximately one third of sequenced species (11, 12). In archaea, selenoprotein utilization was found to date in only three phylogenetic orders (13, 14). Within eukaryotes, selenoproteins are found in most metazoans (including all vertebrates), in some protists and various algae (15, 16), but they are absent in several lineages. In particular, selenoproteinless groups include various orders of insects (17), some nematodes (18), all land plants (but not green algae) (15, 19), and a number of protist lineages (11).

Notably, fungi were also described as a selenoproteinless lineage (11, 15, 19, 20), and were thought to be the only kingdom of life to be entirely devoid of Sec. This was ascribed to a single event of Sec extinction, occurred prior to the radiation of the last common fungal ancestor. In this study, we challenge this notion by showing conclusive genomic evidence for the use of Sec in fungal species belonging to three different phyla: Chytridiomycota, Zoopagomycota and Mucoromycota. Our phylogenetic analysis supports the scenario of vertical inheritance of Sec utilization within eukaryotes, followed by multiple independent events of Sec extinction in fungal lineages. Notably, Sec was lost at the root of Dikarya, resulting in the absence of Sec from the best studied species of this kingdom (including Saccharomyces cerevisiae).

## MATERIAL AND METHODS

### Sequence data

We downloaded all NCBI genome assemblies available for fungi (Supplementary Table T1) using the script ncbi_assembly.py (https://github.com/marco-mariotti/ncbi_db), which wraps the Entrez Bio Python module. When multiple assemblies were available for the same species, only the most recent one was used. This resulted in a set of 1197 fungal genomes that were analysed for the presence of Sec machinery and selenoproteins.

### Gene prediction

Protein-coding gene prediction was performed using Selenoprofiles, a homology-based gene finder developed specifically to identify selenoproteins and related proteins (21, 22). tRNA^Sec^ was searched using the newly developed *ad-hoc* tool Secmarker (23). We initially scanned all fungal genome assemblies for Sec machinery. Then, we searched for all known selenoprotein families in those species with tRNA^Sec^. This resulted in a number of putative genes, both for Sec machinery and selenoproteins, which were then subjected to phylogenetic analysis and manual inspection. Additionally, we searched these fungal genomes for the occurrence of novel selenoprotein families using Seblastian (24, 25), but all such candidates were discarded after manual inspection (data not shown). SECIS elements were searched and plotted using SECISearch3 (24).

### Phylogenetic analysis

In order to provide phylogenetic context, each candidate protein was searched with blastp (26) against the NCBI NR database (27), and the most similar proteins (e-value<1e-5) were downloaded and aligned to the fungal candidates. When more than 300 proteins were matched in NR for a certain family, these were aligned with mafft (28), and 300 sequence representative were selected using Trimal (29). Phylogenetic reconstruction was then performed using the “build” command of the ETE3 package (30), using the workflow “PhylomeDB”. This workflow (31) comprises the use of various aligner programs combined to a consensus alignment with M-Coffee (32). This is then trimmed to remove uninformative and confounding positions using Trimal (29). Neighbor joining phylogenetic reconstruction is then performed to assess the likelihood of 5 different evolutionary models. Lastly, the final tree is computed by maximum likelihood with PhyML (33) using the best evolutionary model. These trees were plotted using custom ETE3 (30) scripts. The descriptions of NR proteins were then inspected to assign protein families to each phylogenetic cluster and annotate them in the tree figures.

### Species phylogeny

For most analysis (Fig. 1, Supplementary Material S1), NCBI taxonomy was used as rough phylogenetic backbone of fungal species. Attempting to trace the events of Sec loss, we then employed a fully resolved tree of fungi including early branching lineages (34). However, various species, whose genome we analysed in this study, were missing from this tree. The tree was thus complemented with NCBI taxonomy information, and manually merged to produce a partially resolved reference tree (Fig. 2). In this process, a number of species were dropped or compressed since non-informative.

**Figure 1.**
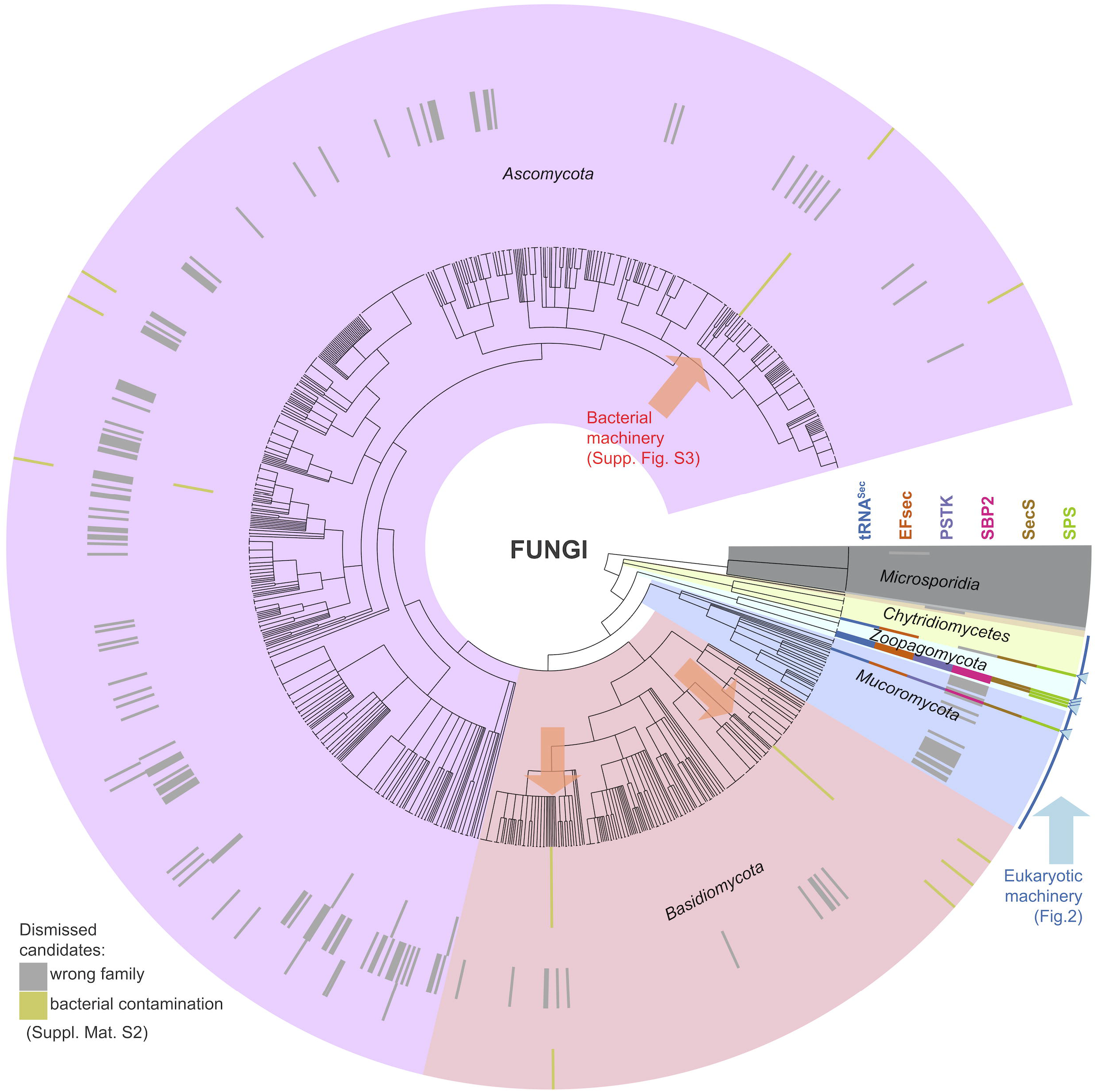
Sec machinery genes in fungal genome assemblies. The figure shows the results of genomic searches for Sec machinery genes (tRNA^Sec^, EFsec, PSTK, SBP2, SecS, SPS) in 1197 species of fungi with publicly available genomes. The tree in the center shows the phylogenetic relationships of species according to NCBI taxonomy, while gene presence predicted by Selenoprofiles is indicated by colored rectangles in the outermost section. Genes in grey and khaki (legend at bottom left) were dismissed by our phylogenetic analysis (Supplementary Material S2) as belonging to a distinct family, or attributed to bacterial contamination, respectively. Colored arrows highlight those genomes containing tRNA^Sec^ genes, divided in two classes (bacterial or eukaryotic) as described in Results. An extended version of this figure including species names is available as Supplementary Figure S1.

**Figure 2.**
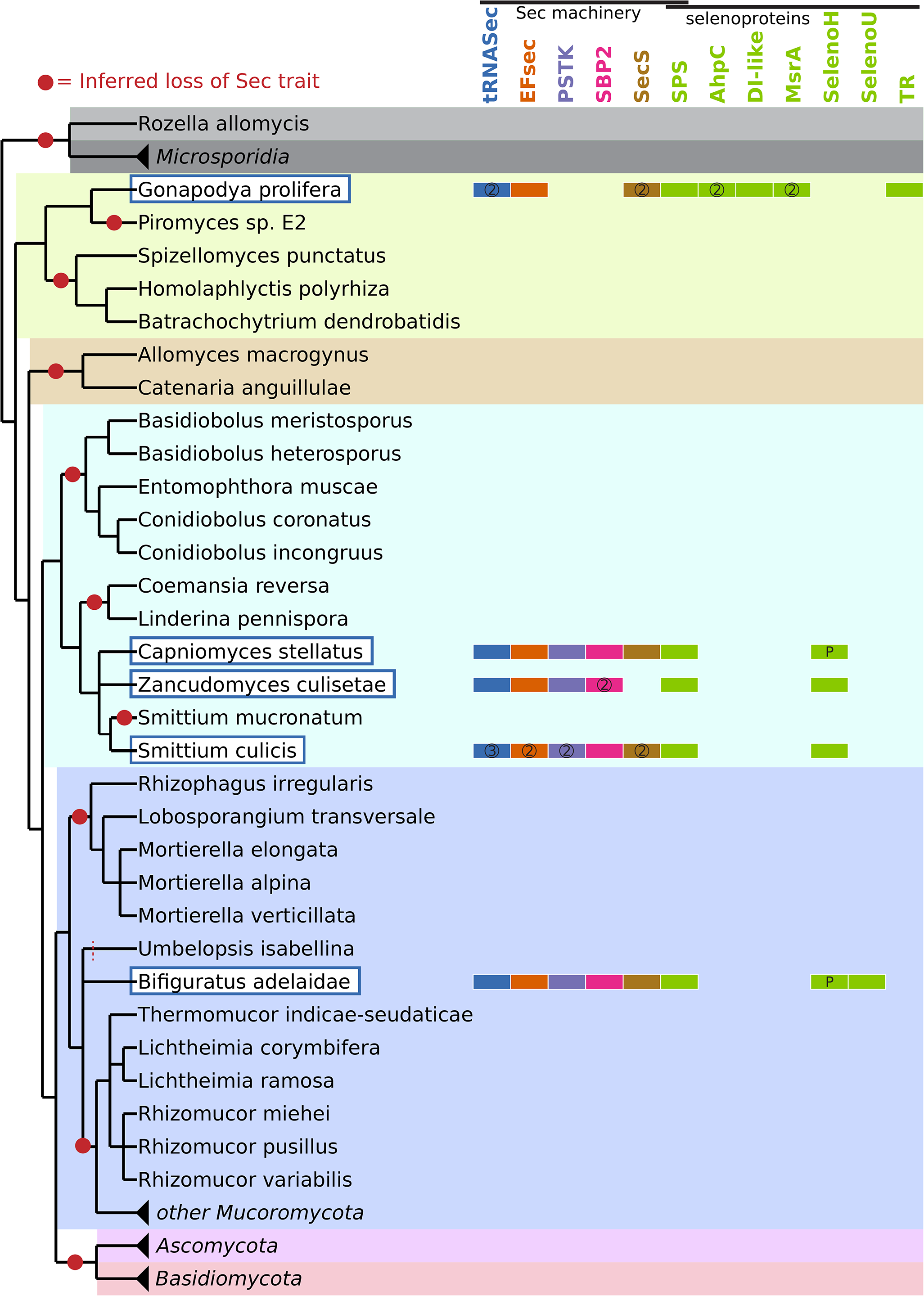
Selenoproteins and Sec machinery in Sec-utilizing fungi. On the left, the figure contains a reference tree of fungi analyzed in this study (Methods). Each species is annotated on the right with the Sec machinery and selenoprotein genes found in their genome assembly. Note that the genes dismissed through phylogenetic analysis are not shown here. Multiple occurrences for a gene family are indicated with circled numbers, while the letter “P” indicates partial gene structures. Sec-utilizing fungi species are highlighted in the reference tree with a blue box and white background. Losses of Sec encoding capacity, inferred by maximum parsimony, are also shown as red circles in the tree. Microsporidia, Ascomycota, Basidiomycota and some Mucoromycota were compressed in this figure for their lack of informative value.

### Phylogenetic profiling

We designed a custom procedure of phylogenetic profiling (35) to identify fungal proteins involved in the Sec pathway. Aiming to detect proteins present in Sec-utilizing genomes but absent in selenoproteinless species, we searched with tblastn the NCBI proteome of Sec-utilizing *Smittium culicis* in all non-dikarya, non-microsporidia fungal genomes, and analysed the e-values of the best matches in each genome. For each query protein, we computed a gene presence threshold based on e-values in Sec-utilizing species, and required e-values worse than such threshold in >90% of non-Sec species. The threshold was computed as the second worst e-value in Sec-utilizing species (thus allowing one Sec-utilizing species to lack candidate proteins), and was further adjusted 20% on log scale (e.g. 1e-10 becomes 1e-8) for higher stringency. We then analysed the resulting candidates to assess the possibility that some constituted novel selenoproteins. We built alignments for each homologous group, and searched them in Sec-utilizing fungal genomes using Selenoprofiles. Other than known selenoproteins in our list, the search did not report any additional in-frame UGA-containing genes. However, selenoprotein genes are typically mis-annotated by standard pipelines, with the Sec codon located either downstream of the annotated gene structure, or downstream of it, or within an artefactually predicted intron (21). By relying on annotated proteins, our profile alignments may have included their deficiencies. We thus considered the possibility that potential Sec residues resided in the immediate proximity of our gene predictions. Separately for upstream, downstream and intron sequences, we considered those regions containing in-frame UGA, but not any other stop codon. Introns were further filtered to require a length multiple of 3. Finally, we translated the regions passing these filters to amino acids, and aligned and manually inspected sequences across species in order to assess their conservation. Based on lack of conservation around all potential Sec-UGAs, all candidates could be dismissed.

## RESULTS

We downloaded all available fungal genomes from NCBI (1197 species, Supplementary Table T1), and examined them for the presence of eukaryotic Sec machinery genes (see Methods) using Selenoprofiles (21, 22) and Secmarker (23). These automatically generated predictions were then analyzed to assess two potential confounders: the presence of protein families sharing similarities with those of interest, and the presence of contaminant sequences in fungal genome assemblies. Both problems can be assessed by a phylogenetic analysis, inspecting the reconstructed gene trees of candidate proteins together with their most similar annotated sequences (see Methods). Our initial genomic predictions showed the presence of Sec machinery genes in a sparse fashion across a number of species (Fig. 1, Supplementary Material S1). However, after our phylogenetic filter (Supplementary Material S2), *bona-fide* Sec machinery proteins were detected only in a handful of genomes, and co-occurred with tRNA^Sec^ (Fig. 1). Therefore, we further focused on those genomes containing tRNA^Sec^, which comprised eight species belonging to diverse taxonomic groups. These species could be clearly divided into two categories, based on the most similar matches to the Sec machinery: three species presented bacterial machinery, while five other species exhibited eukaryotic machinery genes.

### Scattered presence of bacterial Sec machinery and selenoproteins among Dikarya

The genome assemblies of Dikarya species *Puccinia arachidis* (Basidiomycota; Pucciniomycotina), *Clavaria fumosa* (Basidiomycota; Agaricomycotina), and *Beauveria bassiana* (Ascomycota; Pezizomycotina) presented a bacterial-fold tRNA^Sec^ (SelC), together with a bacterial EFsec (SelB) and bacterial SPS (SelD) (Supplementary Material S3). SPS is a selenoprotein family itself, but the genes found in these species had Cys in place of Sec (as previously observed in other organisms (11)). In *C. fumosa*, EFsec and tRNA^Sec^ co-occurred adjacent in the same sequence contig. SecS candidates were predicted in *P. arachidis*, but were dismissed after phylogenetic analysis (Supplementary Material S2). We searched these genomes for selenoprotein genes belonging to known families, resulting in putative selenoprotein genes in each of these genomes. Most of candidate selenoproteins belonged to the FdhA family (formate dehydrogenase A). This is the most widespread selenoprotein in bacteria (36), which has never been observed in eukaryotic genomes. Within these FdhA genes, we identified a structure overlapping the putative Sec-UGA codon, which fitted the bacterial SECIS consensus (37), whereas no eukaryotic SECIS elements (24) were found. The other two selenoprotein predictions had much weaker confidence scores (Selenoprofiles AWSI z-score). One matched the HdrA profile (heterodisulfide reductase A), a selenoprotein family observed only in bacteria and archaea. It was characterized by poor homology with known HdrA, and did not show any SECIS structure. Phylogenetic analyses showed that the closest annotated sequences actually belonged to a different family (electron transferring flavoprotein ubiquinone oxidoreductase; Supplementary Material S2). The last candidate resulted from a match to the TR family (thioredoxin reductase, belonging to the pyridine-nucleotide thiol disulfide oxidoreductase superfamily). This prediction was also characterized by poor sequence identity, and it lacked SECIS. Phylogenetic analyses further showed that this sequence actually matched more closely bacterial glutathione reductase, another enzyme in the same superfamily, rather than TR (Supplementary Material S2). Thus, we concluded that these HdrA and TR selenoprotein candidates were false positives. In contrast, our phylogenetic analysis confirmed the protein family identity of the FdhA, EFsec and SPS candidates in these three Dikarya species. However, these candidates were clearly of bacterial origin, and were scattered across the protein tree (Supplementary Material S2), indicating that the most similar annotated sequences were from different species in each case.

Altogether, our analysis indicates that the presence of Sec machinery and selenoproteins in the *P. arachidis*, *C. fumosa*, and *B. bassiana* genome assemblies can be best explained by sequence contamination from bacteria, and that these fungi do not possess the Sec trait. Although the possibility that these species actually acquired the Sec utilization trait by horizontal gene transfer (HGT) from bacteria cannot be completely excluded, this scenario is extremely unlikely for several reasons. First, protein translation differs substantially between eukaryotes and bacteria, so that it seems improbable that bacterial Sec machinery could be readily “plugged” into a fungal ribosome. Second, none of the Sec machinery and selenoprotein candidates in these genomes have introns; generally, introns are rapidly acquired by genes truly transferred from bacteria to fungi (38). Third, nearly half of these genes matched annotated bacterial homologs with sequence identity >90%, which is also indicative of contamination. Lastly, these three fungal species are only distantly related, and none of their closely related organisms exhibit any traces of the Sec trait. This implies that, if HGT truly occurred, it would have happened so recently that it is limited to single species, and nevertheless occurred in three independent instances. Due to the common incidence of sequence contaminations in public data and the large number of genomes analyzed in our study, we considered that a prerequisite to make inferences was the observation of the same pattern in multiple species, and the consistency of their phylogenetic signal. For these reasons, we dismissed the Sec-encoding candidate species in Dikarya.

### Eukaryotic Sec machinery and selenoproteins in five non-Dikarya fungal species

We identified eukaryotic Sec machinery (Fig. 2) in the fungal species *Bifiguratus adelaidae* (Mucoromycota; Mucoromycotina), *Gonapodya prolifera* (Chytridiomycota; Monoblepharidomycetes), *Capniomyces stellatus*, *Zancudomyces culisetae* and *Smittium culicis* (Zoopagomycota; Kickxellomycotina). These five species are further referred to as the Sec-utilizing fungi. We identified selenoproteins in each of these genomes, belonging to seven known selenoprotein families. One of them, SPS (Fig. 3), was found as selenoprotein in all Sec-utilizing fungi. SPS is a part of the Sec machinery, as well as selenoprotein enzyme itself in many species across prokaryotes and eukaryotes (11). *G. prolifera* was the species with the largest number of selenoproteins (Fig. 2). Besides SPS, these included AhpC (alkyl hydroperoxide reductase C), an oxidoreductase previously identified as selenoprotein in certain bacteria, protists and porifera (24); MsrA (methionine sulfoxide reductase A) (39), previously identified as selenoprotein in algae, protists, and various non-vertebrate metazoa (40); DI-like, a protein homologous to vertebrate iodothyronine deiodinases (41), and present as selenoprotein also in various invertebrates, protists and bacteria; and finally TR, found as a selenoprotein across most Sec-utilizing eukaryotic lineages. Notably, the Sec-containing TR in *G. prolifera* constitutes the first case of animal-like TR described in fungi, since species in this kingdom possess a distinct, shorter and Sec-independent form of TR (42). Two additional selenoprotein families were found in Sec-utilizing fungi that were missing in *G. prolifera* (Fig. 2): SelenoU, a selenoprotein in many eukaryotes including fish, which was converted to Cys homolog in therian mammals (9); and SelenoH, found as selenoprotein in mammals and other metazoa including insects (43). SelenoH was well conserved in Sec-utilizing fungi: it was found in all genomes except *G. prolifera*, wherein, nevertheless, we detected a gene fragment similar to its C-terminal domain (data not shown).

**Figure 3.**
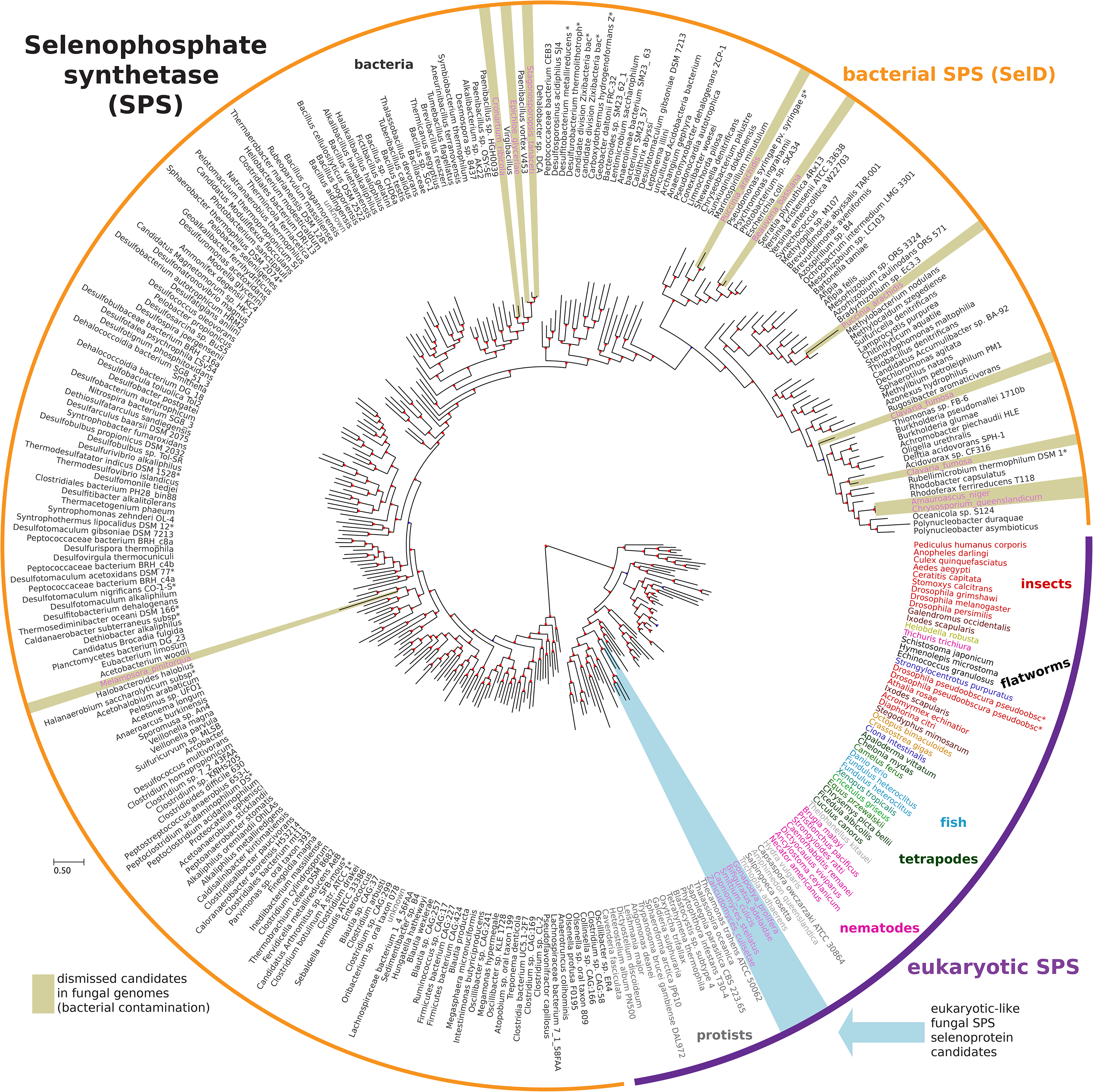
Reconstructed phylogenetic tree of SPS proteins. This tree was built based on the sequences of SPS candidates in fungal genomes, aligned with their most similar proteins annotated in NCBI NR. The source species is shown for each protein, and it is colored according to its taxonomic group. Colored arrows highlight the fungal SPS candidates, divided in two classes as described in Results. Analogous trees for the rest of Sec machinery and selenoprotein genes are available in Supplementary Material S2.

We examined these selenoprotein genes for the presence of eukaryotic SECIS elements. Surprisingly, we found a SECIS element only in 5 out of the 16 fungal selenoprotein candidates (Fig. 4), even when setting the SECISearch3 parameters for maximum sensitivity (24). The identified fungal SECIS elements fit the eukaryotic consensus. Two of them, belonging to *G. prolifera* AhpC genes, lacked the typically conserved apical adenines, but this feature is not universal in SECIS elements (2). As expected, SECISes were found downstream of the coding sequences of selenoprotein genes, starting at a distance between ~40 and ~420 nucleotides after the stop codon.

**Figure 4.**
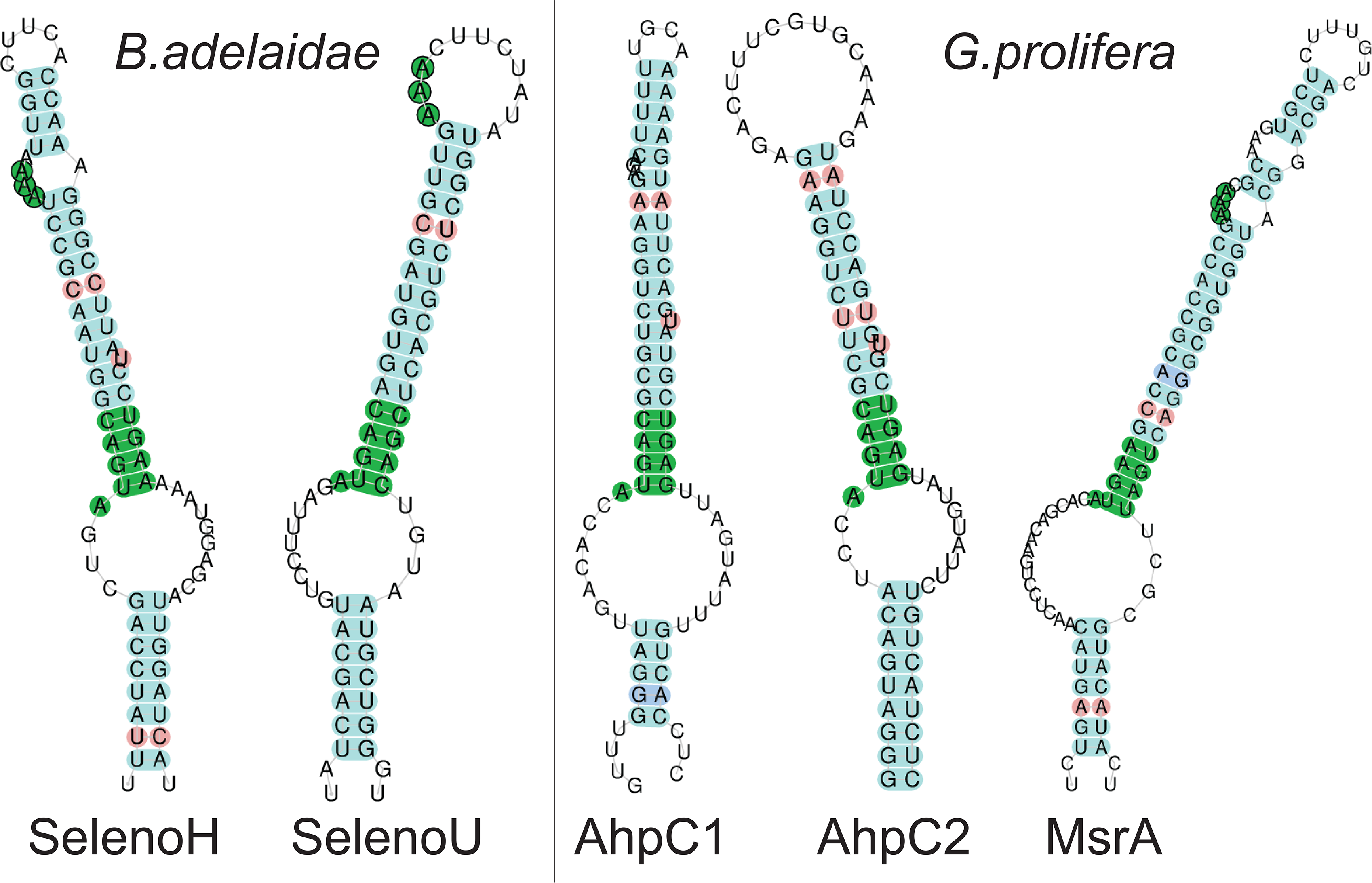
Eukaryotic SECIS elements identified in fungal selenoproteins. These SECIS elements were identified downstream of the coding sequences of some of the fungal selenoprotein genes (24). Selenoproteins not listed here (but shown in Fig. 2) could not be assigned any SECIS elemen

Despite the lack of identifiable SECIS in the rest of selenoprotein candidates, we are confident that they represent bona-fide selenoproteins based on the fact that they show homology to known selenoprotein families, and possess a TGA codon aligned precisely with the known Sec position. All candidates passed a manual inspection, and our phylogenetic analysis (Methods, Supplementary Material S2) confirmed their protein family identity. In turn, this suggests that fungi may use distinct SECIS element variants undetected by current methods of SECIS prediction.

Attempting to identify potential novel selenoproteins in the genomes of Sec-encoding fungi, we also applied the Seblastian pipeline (24), which uses SECIS identification as a first step. Seblastian successfully predicted the selenoprotein genes with identifiable SECIS reported above (Fig. 4), and reported additional low scoring candidates for novel families. All these candidates could be discarded after manual analysis (data not shown).

### Vertical inheritance of Sec machinery within eukaryotes

The phylogenetic analysis of Sec machinery proteins gave a strong support to the hypothesis that *B. adelaidae*, *G. prolifera*, *C. stellatus*, *Z. culisetae* and *S. culicis* are true Sec-encoding fungi. Importantly, the Sec machinery protein trees recapitulate the species tree, which is consistent with the vertical inheritance of the trait. These results are illustrated for the SPS family in Fig. 3, while the rest of protein trees are provided in Supplementary Material S2. In the reconstructed protein tree, SPS sequences from these species form a monophyletic cluster, as expected according to their species phylogeny. This supports the scenario of common descent, with the Sec-containing SPS gene inherited from the ancestor of fungi to these extant species. The same clustering pattern is observed also for PSTK and SBP2, while fungal EFsec and SecS form instead two clusters. The position of the fungal clusters relative to the homologs in other lineages is also informative: they are located in the expected position as outgroup of metazoa, sometimes within protist lineages. Beyond fungal species, Sec machinery trees are in general accordance with the known species phylogeny, with some exceptions. In particular, nematodes and insects are placed in trees in a more basal position than expected, located among protist groups. This may be caused by an acceleration of sequence divergence in this lineage, causing long-branch attraction in phylogenetic reconstruction.

In a previous study (11), we concluded that the Sec encoding trait has been likely directly inherited from the root of eukaryotes to extant Sec-utilizing species, while the various non-Sec-utilizing groups underwent a number of parallel Sec losses. The phylogenetic trees presented here are consistent with this scenario, and thus support direct and continuous inheritance of the Sec trait from the root of eukaryotes to Sec-utilizing fungi.

### Putative genes related to Sec trait predicted by phylogenetic profiling

Sec utilization appears sparse across early branching fungal lineages, suggesting a highly dynamic evolutionary process. Given the remarkable diversity spanned by fungal genomes, we hypothesized that phylogenetic profiling could be profitably taken to discover fungal genes related to the Sec trait. This technique (35, 44) exploits gene co-occurrence across genomes to link genes to pathways. Aiming to uncover potential novel selenoproteins and proteins involved in the Sec pathway, we developed a custom phylogenetic profiling procedure (see Methods) to identify genes present in Sec-utilizing fungal genomes but absent in selenoproteinless species. Our procedure resulted in a list of 62 candidate proteins clustered in 27 homologous groups (Supplementary Table T2). Proteins PSTK, EFsec, SecS and selenoproteins SPS and SelenoH emerged as top scoring candidates, supporting robustness of the procedure. The rest of candidates had diverse annotated functions, and included oxidoreductases, transporters, uncharacterized proteins and others. We further examined candidates as potential novel selenoproteins, seeking conserved Sec-UGA codons, but found none (Methods). These genes may be involved in Sec transport, metabolism, or regulation, or just represent false positives. Future experiments will be necessary to validate candidates and clarify their role in Sec biology.

## DISCUSSION

In this study, we report that Sec is genetically encoded by several species of fungi, defying the paradigm that this kingdom of life uses only the 20 canonical amino acids. Through genomic searches and sequence analyses, we show that *B. adelaidae*, *G. prolifera*, *C. stellatus*, *Z. culisetae* and *S. culicis* possess the genetic machinery necessary to synthesize, encode and decode Sec. We also identified several selenoprotein genes in these species (Fig. 2). Techniques of phylogenetic reconstruction, applied to the fungal gene candidates aligned with their most similar annotated proteins, allowed us to differentiate between artefactual gene occurrences and well supported cases of Sec-encoding fungal genomes. In particular, our analyses led us to dismiss the Dikarya species *P. arachidis*, *C. fumosa*, and *B. bassiana* as Sec encoding organisms. Their available genome assemblies exhibit traces of Sec machinery, but we attributed them to bacterial contaminations of the assemblies. In contrast, Sec-utilizing fungi possess genuine eukaryotic Sec machinery and selenoprotein genes whose phylogenetic signal is consistent with their taxonomy, placing them in most cases in a single monophyletic cluster in reconstructed protein trees.

Our analyses indicate that, in all likelihood, the Sec trait was vertically inherited from early eukaryotes to fungi; subsequently, many fungal lineages lost Sec in multiple independent events. There are two alternative scenarios that could potentially explain the observed pattern of gene presence. First, Sec machinery and selenoproteins in the genome assemblies of Sec-utilizing fungi may derive from sequence contamination. Given the results of our phylogenetic analyses, such contaminating sequences would necessarily belong to the same organism (or a closely related group) in all cases, and constitute a Sec-containing eukaryotic lineage with phylogenetic signal indistinguishable from that of fungi. However, the genomes of Sec-utilizing fungi were generated by three different research organizations (focusing on different fungal phyla), so that the transversal occurrence of the same contamination seems highly unlikely. The second alternative scenario is that, in one or more cases along the fungal tree, HGT may have occurred to transfer the Sec trait from a Sec-containing eukaryote to fungi. This scenario seems even less likely: Sec-utilizing lineages span very diverse niches, and exposure to the same HGT donor appears implausible. Furthermore, while Sec machinery genes are sometimes clustered in operon-like structures in bacteria, they are scattered throughout chromosomes in eukaryotic genomes, so that multiple subsequent HGT events would be required to transfer a functional Sec trait. We conclude that the vertical and continuous inheritance of the Sec trait is the most reasonable scenario.

We thus predict that Sec machinery was never lost by Sec-utilizing fungi. It is conceivable, nevertheless, that some of their extant selenoproteins were brought by horizontal transfer. A prime candidate for such an event is the Sec-containing TR in *G. prolifera*. In fact, this gene is the only large (animal-like) TR found in the fungal kingdom. These organisms, together with plants, possess another unrelated, shorter TR (42). There are two possible evolutionary scenarios to explain this pattern of occurrence: either the animal-like TR family was lost independently in many lineages concomitant with the Sec trait, or it was lost early in fungal phylogeny, functionally replaced by a short non-Sec TR, and then *G. prolifera* acquired a Sec-TR by HGT from a Sec-utilizing protist (based on the reconstructed TR tree, Supplementary Material S2). Current data does not allow to resolve between these scenarios.

Based on sequence homology and phylogenetic signal, we are confident that all our selenoprotein candidates in these organisms constitute *bona-fide* cases of Sec encoding genes. However, we were surprised to find that many sequences lacked recognizable SECIS elements. Although we do not yet know why this is the case, we speculate that this may be related to the high degree of divergence that we observe for SBP2, their partner protein (Supplementary Material S2). Sec-utilizing fungi appear to possess a limited number of selenoproteins (although some may remain to be discovered). This may have facilitated co-evolutionary drift of SBP2 together with the functional consensus of SECIS elements. We are confident that future studies will elucidate the identity of fungal SECISes. In particular, the availability of additional genomes of Sec-utilizing fungi will allow the application of comparative strategies for *de novo* prediction of functional RNA structures, which is impractical at the current stage.

The discovery of selenoproteins in fungi may open the way for the production of selenoproteins in bio-engineered yeast. The synthesis of exogenous selenoproteins has previously been attempted in *S. cerevisiae*: all human Sec machinery genes were transferred to yeast and successfully expressed, but Sec insertion was never observed (45). This has been speculatively attributed to incompatibilities between vertebrates and fungi, due to differences in ribosome structure and translation mechanism. Therefore, we believe that the use of Sec machinery native to the fungal world may instead lead to fruitful results.

The Sec-utilizing fungi identified in this study belong to various early-branching phyla: Chytridiomycetes, Zoopagomycota and Mucoromycota. Each of these lineages includes several other species with available genomes, which show no traces of Sec machinery or selenoprotein genes. The scattered distribution of Sec-utilizing species implies many independent losses of the Sec trait in fungi. Even in the most parsimonious scenario (Fig. 2), we infer at least 10 different Sec losses in fungi. One of these losses was mapped prior to the split of Dikarya (Ascomycota and Basidiomycota, diverged around 700 mya (46)), resulting in the lack of Sec in *Saccharomyces cerevisiae* and most other best studied fungi. Such high incidence of parallel losses may seem surprising; however, this was previously well documented for the Sec trait at various evolutionary scales. Parallel Sec losses were frequently observed in prokaryotes (11, 14, 47, 48), in various protist groups (11, 19), and even within animals, with at least 5 independent Sec loss events meticulously traced within insects (17, 40). Interestingly, selenoproteinless insects retained a number of genes that originally were selenoproteins. In these genes, Sec-UGA codons were converted to Cys codons, presumably maintaining the molecular function of the encoded protein products. In fungi, however, we did not observe any Sec-to-Cys conversions: no orthologs were found for selenoprotein genes in selenoproteinless species. This surprising observation suggests an different mechanism for Sec losses in fungi, wherein the utility of the functions of selenoproteins themselves (rather than the advantage provided by Sec over Cys in those proteins) faded and led to their loss (rather than their conversion to Cys).

The observation of so many Sec losses within fungi compels to investigate the selective forces driving Sec maintenance and loss, and any possible commonalities between fungi and other lineages undergoing Sec extinctions. Sec losses constitute an instance of convergent parallel evolution (from a Sec encoding state to Sec devoid). As such, their causes may be rooted in either (i) environmental changes affecting parallel lineages at once (e.g. drastic decrease world-wide in bioavailability of selenium); or (ii) important changes fixed in the period preceding a large evolutionary radiation, later causing a “snowball” effect (e.g. if early fungi evolved a different physiology or lifestyle which diminished their exposure to oxidants, utility of selenoproteins would have been attenuated). Conceivably, future studies will elucidate some of these issues. To that end, the functional characterization of selenoproteins and related genes in Sec-utilizing fungi will be key to clarify why these organisms have clung to them for so long, and, by extension, why so many other species could instead lose all Sec genes with no apparent consequence.

## DATA AVAILABILITY

The latest version of the selenoprotein gene finder software Selenoprofiles is available at https://github.com/marco-mariotti/selenoprofiles. Fungal genome assemblies used in this study (Supplementary Table T1) can be found at NCBI website https://www.ncbi.nlm.nih.gov/assembly, or downloaded in batch using the script ncbi_assembly at https://github.com/marco-mariotti/ncbi_db.

## SUPPLEMENTARY DATA

Supplementary Table T1. Species names, assembly identifiers and taxonomic annotation all of fungal genomes analyzed in this study.

Supplementary Table T2. Results of phylogenetic profiling to detect proteins related to Sec. The table lists all *S.culicis* query proteins predicted as present in all (or all except one) Sec-utilizing fungal genomes, and absent in at least 90% of non-Sec-utilizing fungal genomes (Methods). The e-values of the best tblastn matches are listed for each species, followed by the annotated query description, a manual annotation compiled inspecting the titles of similar proteins at NCBI, and the assignment to a cluster of similar candidates.

Supplementary Figure S1. Sec machinery genes in fungal genome assemblies (extended version). This figure is an extension of Fig. 1 that includes species names, allowing to inspect the identity of each genome analyzed.

Supplementary Material S2. Phylogenetic trees for Sec machinery and selenoprotein analyzed in this study. This document contains phylogenetic trees for the proteins PSTK, SBP2, SecS, EFsec, HdrA, TR, FdhA, MsrA, DI, SelenoH, SelenoU and AhpC. Trees were built, plotted and annotated with same procedure as the SPS tree shown in Fig. 3 (Methods). Note that AhpC and SelenoU were joined in a single tree since they share sequence similarity.

Supplementary Figure S3. Bacterial Sec machinery and selenoproteins predicted in scattered Dikarya genome assemblies. The figure shows the Sec machinery and selenoproteins predicted in *P.arachidis*, *C.fumosa* and *B.bassiana*, the only three Dikarya species with tRNA^Sec^. These species are not monophyletic in our species set, and instead constitute isolated cases among Dikarya (see Fig. 1). Note that the phylogenetic analysis indicated that the SecS, HdrA and TR candidates actually belonged to other protein families, resulting in their dismissal. Taken together, our results indicate that the presence of tRNA^Sec^, EFsec, SPS and FdhA genes in these genomes is due to assembly contamination from various Sec-utilizing bacterial species.

Supplementary Data are available at NAR online.

## ACKNOWLEDGEMENTS

Not applicable.

## FUNDING

This work was supported by the National Institutes of Health grants DK117149 and CA080946. Funding for open access charge: National Institutes of Health.

## CONFLICT OF INTEREST

The authors declare that they have no competing interests.

